# In Silico Design and Evaluation of CRISPR Cytosine Base Editing Strategy for Potential Correction of the ASPA Variant Associated with Canavan Disease

**DOI:** 10.64898/2026.10.24.761534

**Authors:** Aditi Verma, Elamathi Natarajan

**Affiliations:** Department of Bioinformatics, AI & Machine Learning, Biotechnika Info Labs Pvt. Ltd., Bengaluru, Karnataka, India

**Keywords:** CRISPR-Cas9, Canavan Disease, Cytosine base editing, CRISPRitz, ASPA gene, Genome editing

## Abstract

**Background:** Canavan disease is a rare autosomal recessive neurodegenerative leukodystrophy caused by pathogenic variants in the ASPA gene, for which effective molecularly targeted therapies remain limited. The NM_000049.4:c.691T>C (p.Tyr231His) variant introduces a pathogenic C•G base pair that is potentially amenable to correction using cytosine base editing (CBE). This study aimed to design and computationally evaluate a CBE-based CRISPR strategy for potential correction of this ASPA variant.

**Methods:** An integrated in silico workflow was used to characterize the ASPA p.Tyr231His variant and design a CBE-compatible sgRNA. Variant pathogenicity, conservation, and structural stability were assessed using multiple computational tools. Candidate sgRNAs were designed and evaluated for editing-window positioning, PAM compatibility, specificity, GC content, bystander editing, and secondary structure. Genome-wide off-target potential was further assessed using CRISPRitz against the GRCh38 reference genome.

**Results:** The ASPA p.Tyr231His variant was exceptionally rare (gnomAD allele frequency: 1.86 × 10^−6^), highly conserved, and consistently predicted to be deleterious, with DynaMut2 indicating reduced protein stability (ΔΔG = −2.04 kcal/mol). Of two candidate sgRNAs, one was selected based on optimal editing-window positioning, absence of predicted bystander editing, favourable GC content (45%), and specificity scores (MIT: 82; CFD: 93). RNAfold predicted minimal secondary-structure constraints (MFE: 0.00 kcal/mol). CRISPRitz identified 108 potential off-target sites, with none containing fewer than three mismatches, no bulge-associated sites, and only three predicted sites within exonic regions.

**Conclusions:** The integrated computational analysis identified a candidate sgRNA with favourable predicted editing characteristics and genome-wide specificity for potential CBE-mediated correction of the ASPA c.691T>C (p.Tyr231His) variant. These findings provide a rational framework for prioritizing this guide for experimental evaluation. Further cell-based and in vivo studies are required to establish editing efficiency, editing purity, bystander activity, off-target effects, and restoration of ASPA function before therapeutic applicability can be assessed.

## 1. Introduction

Gene-editing technologies offer a promising strategy for correcting disease-causing genetic mutations at their source. Their therapeutic potential is especially significant for hereditary monogenic disorders, which result from pathogenic variants in a single gene (Roth and Marson 2021). CRISPR-Cas9 genome editing has transformed biomedical research by enabling precise and efficient modification of genomic sequences. This technology holds significant promise for the treatment of genetic disorders, many of which are caused by single-nucleotide mutations. However, despite its versatility, the clinical application of CRISPR-based systems is constrained by challenges such as off-target effects and unintended genomic alterations, which raise concerns regarding safety and specificity. Advanced computational techniques are increasingly being used to optimize CRISPR systems and solve these problems (Abudayyeh and Gootenberg 2024).

Leukodystrophies, sometimes referred to as myelin or white matter (WM) diseases, are caused by downstream metabolic divergence in neuronal and/or non-neuronal populations or functional mutations in glia (Seitelberger 1984; Powers and Rubio 1995; Kevelam et al. 2016). Among these disorders, Canavan disease is a rare autosomal recessive leukodystrophy caused by pathogenic variants in the ASPA gene, which is predominantly expressed in mature oligodendrocytes of the central nervous system. ASPA expression is predominantly observed in mature oligodendrocytes of the central nervous system, where its deficiency contributes to impaired myelin maintenance and the development of Canavan disease (Kirmani et al. 2002; Moffett et al. 2007).

Aspartoacylase (UniProt ID: P45381 · ACY2_HUMAN) (ASPA) is a 313-amino acid metabolic enzyme that plays an important role in central nervous system physiology by catalysing the conversion of N-acetyl-L-aspartate (NAA) into acetate and aspartate (Birnbaum et al. 1952; Birnbaum 1955; Kaul et al. 1993). ASPA activity is particularly relevant in oligodendrocytes, where regulation of NAA metabolism contributes to normal myelin development and maintenance (Kirmani et al. 2002, 2003; Sommer and Sass 2012). Pathogenic variants in the ASPA gene disrupt enzyme function and lead to accumulation of NAA, resulting in impaired oligodendrocyte activity and defective myelination (Traka et al. 2008). These molecular abnormalities underlie Canavan disease, an autosomal recessive neurodegenerative leukodystrophy characterized by progressive neurological impairment. Therefore, restoration of functional ASPA through precise correction of disease-associated nucleotide changes represents a promising therapeutic strategy.

CRISPR-Cas9 outperforms previous engineered nucleases, such as zinc-finger nucleases and transcription activatorlike effector nucleases, in terms of simplicity and flexibility by using a single effector protein and a configurable guide RNA (gRNA) (Gupta and Musunuru 2014). The PAM, a brief conserved DNA sequence located directly downstream of the target site, further determines the exact targeting of Cas9 (Vink et al. 2021). This sequence is usually 5′-NGG3′ for SpCas9, where “N” can be any nucleotide (Miller et al. 2020). Poorly designed sgRNAs may lead to significant off-target effects, potentially causing unintended mutations and limiting therapeutic applicability. Improvements in gRNA design and delivery have enhanced both target coverage and precision. For instance, designing focused libraries that target druggable gene families has improved the efficiency of screening experiments. Similarly, combining CRISPR screening with drug-based treatments has helped uncover subtle chemogenetic relationships by revealing drug–gene synergies through patterns of guide RNA depletion data. (Joung et al. 2017; Tsiami et al. 2024).

In this study, we aimed to design and evaluate a highly specific sgRNA for cytosine base editing targeting a pathogenic mutation in the ASPA gene associated with Canavan disease. An integrated computational pipeline was employed, incorporating multiple bioinformatics tools to assess sgRNA efficiency, structural stability, and genome-wide specificity. This approach provides a systematic framework for optimizing CRISPR-based therapeutic strategies and highlights the potential of in silico methodologies in precision genome editing.

## 2. Materials and Methods

### 2.1 Sequence retrieval and mutation identification

The ASPA transcript sequence (NM_000049.4) was retrieved from the NCBI RefSeq database, while pathogenic variant information was obtained from ClinVar (Rady et al. 1999; Zano et al. 2013). The ASPA gene is located on chromosome 17p13.2, and genomic coordinates were determined according to the GRCh38 human genome assembly. The pathogenic missense variant NM_000049.4:c.691T>C (p.Tyr231His) was selected for CRISPR-based therapeutic evaluation. This variant results from a thymine-to-cytosine substitution at coding nucleotide position 691, replacing the conserved tyrosine residue with histidine at amino acid 231. The selected variant was subsequently evaluated for its suitability for correction using a cytosine base editing approach.

### 2.2 Variant annotation and in silico analysis

The identified ASPA gene variant was further characterized through comprehensive annotation and computational analysis. To assess the population frequency and rarity of the mutation, allele frequency data were retrieved from the Genome Aggregation Database (gnomAD v4.1.1) (https://gnomad.broadinstitute.org/) (Karczewski et al. 2020). Additionally, the functional impact of the ASPA c.691T>C (p.Tyr231His) variant was evaluated using MutationTaster2 (https://www.mutationtaster.org/) (Schwarz et al. 2014) based on the GRCh38 human genome assembly.

MutationTaster was used to predict the potential functional consequences of the nucleotide substitution and classify the variant according to its predicted effect on gene function. Additional pathogenicity scores including CADD, REVEL, PolyPhen-2, and SIFT were retrieved from publicly available variant annotation resources and used for comparative functional assessment.

Evolutionary conservation of Tyr231 was assessed by performing multiple sequence alignment of ASPA protein sequences from human, mouse, rat, Xenopus, and zebrafish using Clustal Omega (https://www.ebi.ac.uk/jdispatcher/msa/clustalo) (Madeira et al. 2024). The resulting alignment was analysed to determine whether the affected residue was conserved across vertebrate species, providing additional evidence regarding its functional importance. Collectively, these analyses established the rarity, predicted pathogenicity, and evolutionary conservation of the ASPA p.Tyr231His variant prior to downstream structural and genome-editing analyses.

### 2.3 Structural modelling and stability analysis of the ASPA p.Tyr231His variant

Protein structural prediction confidence assessment was performed using AlphaFold2 through the Google Colab implementation (Jumper et al. 2021; Mirdita et al. 2022). The ASPA amino acid sequence containing the p.Tyr231His variant was submitted to AlphaFold2, and prediction reliability was evaluated using multiple sequence alignment (MSA) coverage and predicted Local Distance Difference Test (pLDDT) scores. The pLDDT scores provided residuelevel confidence estimates, while the MSA coverage plot was used to assess the availability of homologous sequence information supporting the prediction. These parameters were used to evaluate the reliability of the AlphaFold2 prediction prior to downstream structural stability analysis.

To further investigate the effect of the p.Tyr231His substitution on protein stability, the experimentally determined human ASPA crystal structure (PDB ID: 2O53) was analysed using the DynaMut2 web server (https://biosig.lab.uq.edu.au/dynamut2/) (Rodrigues et al. 2021). The mutation was introduced into chain A, and DynaMut2 was used to predict the change in folding free energy (ΔΔG) associated with the amino acid substitution, thereby estimating the mutation-induced effect on protein stability.

### 2.4 Base editor selection and sgRNA design for CRISPR-mediated editing

The reference sequence of the ASPA gene was used as input for guide RNA design. The target site corresponding to the pathogenic c.691T>C (p.Tyr231His) variant was identified, and candidate single-guide RNAs (sgRNAs) were designed using the BE-Designer web server (http://www.rgenome.net/be-designer/) (Hwang et al. 2018). The platform was used to identify candidate protospacer sequences, protospacer-adjacent motif (PAM) sites, GC content, and the position of the target nucleotide within the base-editing window. Candidate guides were prioritised based on the presence of a canonical NGG PAM, appropriate GC content (40–60%), correct positioning of the pathogenic cytosine within the optimal editing window, and the absence of predicted bystander editing.

The ASPA c.691T>C variant results in conversion of the wild-type tyrosine codon (TAC) to the pathogenic histidine codon (CAC). This substitution introduces a pathogenic C•G base pair that can potentially be corrected through cytosine base editing by converting C•G to T•A, thereby resulting in conversion of the mutant CAC codon towards the wild-type TAC codon. Therefore, a cytosine base editor was selected for evaluation of this variant. CBEs catalyse the deamination of cytosine to uracil within a defined editing window, enabling C•G-to-T•A conversion without generating double-strand DNA breaks or requiring donor DNA templates.

sgRNA design was performed in BE-Designer using the YE1/EE/YEE-BE (C→T) base editor configuration. The nucleotide corresponding to the ASPA c.691T>C variant was positioned at the sixth nucleotide of the protospacer, within the optimal editing window (positions 4–8), enabling potential C→T correction while minimizing predicted bystander editing of neighbouring cytosines.

### 2.5 Evaluation of candidate sgRNAs

Evaluation of candidate sgRNAs was performed using CRISPOR (https://crispor.gi.ucsc.edu/crispor.py) (Concordet and Haeussler 2018), which provides predictive scoring metrics for guide RNA performance. MIT specificity scores and CFD (Cutting Frequency Determination) scores were used to assess guide specificity and potential off-target activity. sgRNAs exhibiting higher MIT and CFD specificity scores and favourable predicted editing characteristics were selected for downstream analyses. The final sgRNA was selected based on combined evaluation of specificity scores, predicted bystander editing, PAM compatibility, GC content, and editing window positioning.

The secondary structure of the selected sgRNA was predicted using the RNAfold web server from the ViennaRNA package 2.0 (ViennaRNA Web Services) (Lorenz et al. 2011). The sgRNA sequence was analysed to determine minimum free energy (MFE), thermodynamic ensemble properties, ensemble diversity, and predicted secondary structure. RNAfold-generated structural models were subsequently used to evaluate sgRNA folding characteristics and structural accessibility, which are important factors influencing guide RNA stability and target recognition efficiency.

### 2.6 Genome-wide off-target analysis using CRISPRitz

To assess the genome-wide specificity of the designed sgRNA, off-target analysis was performed using CRISPRitz (https://github.com/pinellolab/CRISPRitz) (Cancellieri et al. 2020). The human reference genome (GRCh38/hg38) was used as the alignment database. The sgRNA sequence was searched against the genome to identify potential offtarget sites, allowing up to four mismatches between the guide RNA and genomic target sequences.

To improve detection sensitivity, both mismatch-based and bulge-aware searches were performed to identify potential off-target sites involving sequence mismatches or non-canonical DNA/RNA bulge interactions. Predicted off-target sites were subsequently annotated according to their genomic context, including exonic, intronic, promoter, enhancer, transcription factor binding, and other regulatory regions.

The resulting datasets were used to evaluate mismatch distribution, positional mismatch patterns, genomic distribution of off-target sites, and overall sgRNA specificity. This comprehensive workflow provided a genome-wide assessment of potential off-target activity associated with the designed sgRNA.

## 3. Results

### 3.1 Comprehensive genomic characterization and in silico functional evaluation of the ASPA variant

Population frequency analysis demonstrated that the ASPA p.Tyr231His variant is an exceptionally rare pathogenic allele, with a gnomAD allele frequency of 1.86 × 10^−6^. The substitution replaces tyrosine at amino acid position 231 with histidine, a change predicted to alter the physicochemical properties of the residue.

Sequence coverage analysis demonstrated extensive homologous sequence support across the ASPA protein, indicating strong evolutionary information supporting the AlphaFold2 prediction (Fig. 1a). The predicted Local Distance Difference Test (pLDDT) scores exceeded 90 across the majority of residues, indicating high confidence in the predicted structural regions, with reduced confidence observed only in localized flexible loop and terminal regions (Fig. 1b).

**Fig 1.**
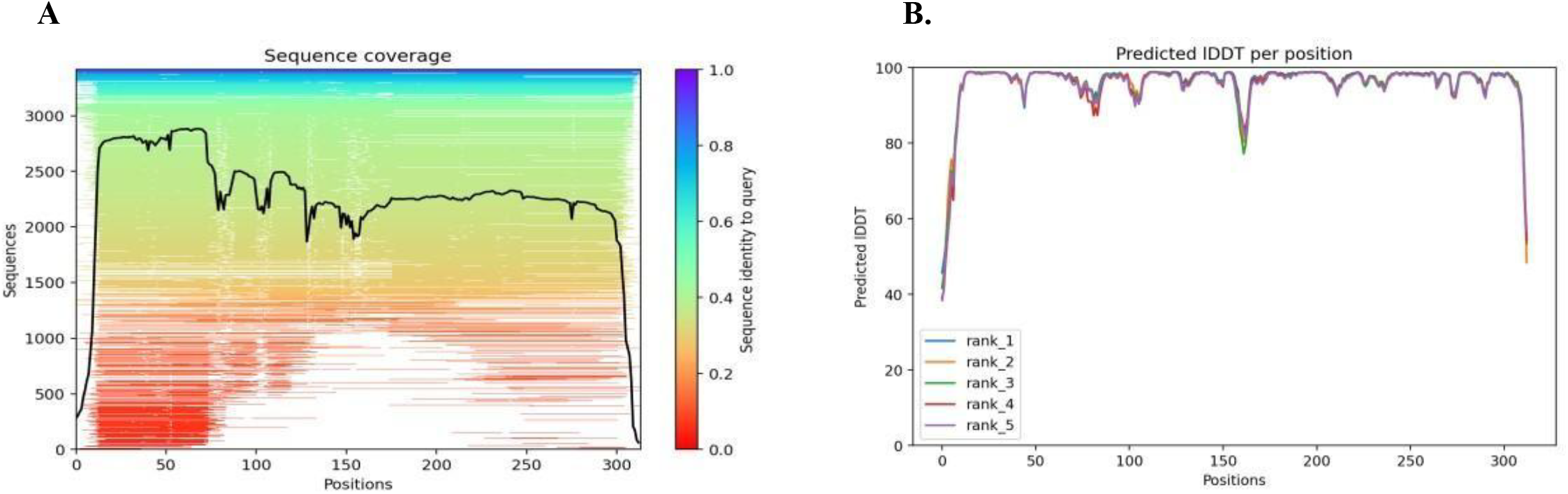
Structural quality assessment of the AlphaFold -predicted ASPA Y231H variant. (A) Sequence coverage plot showing the distribution and depth of homologous sequences across the ASPA protein sequence. The black line represents the number of aligned sequences covering each residue position, while the colour scale indicates sequence identity to the query sequence. (B) Predicted local distance difference test (pLDDT) scores across the ASPA sequence. Higher pLDDT values indicate greater confidence in the predicted protein structure.

DynaMut2 analysis predicted that the Y231H missense mutation exerts a destabilizing effect on the protein, with a calculated ΔΔG value of −2.04 kcal/mol. Structural comparison of the wild-type and mutant proteins revealed that the native Tyr231 residue participates in an extensive network of hydrophobic and aromatic interactions that contribute to local structural stability. The wild-type Tyr231 residue (Fig. 2A) forms an extensive network of hydrophobic and aromatic interactions with neighboring residues, contributing to local structural stability and compact packing. In contrast, the mutant His231 residue (Fig. 2B) exhibits reduced hydrophobic interactions and the formation of alternative polar interactions, indicating alteration of the native interaction pattern. These structural changes are consistent with the predicted decrease in protein stability and suggest that the Y231H substitution may perturb the local conformational environment, potentially affecting local protein stability and conformational dynamics.

**Fig. 2.**
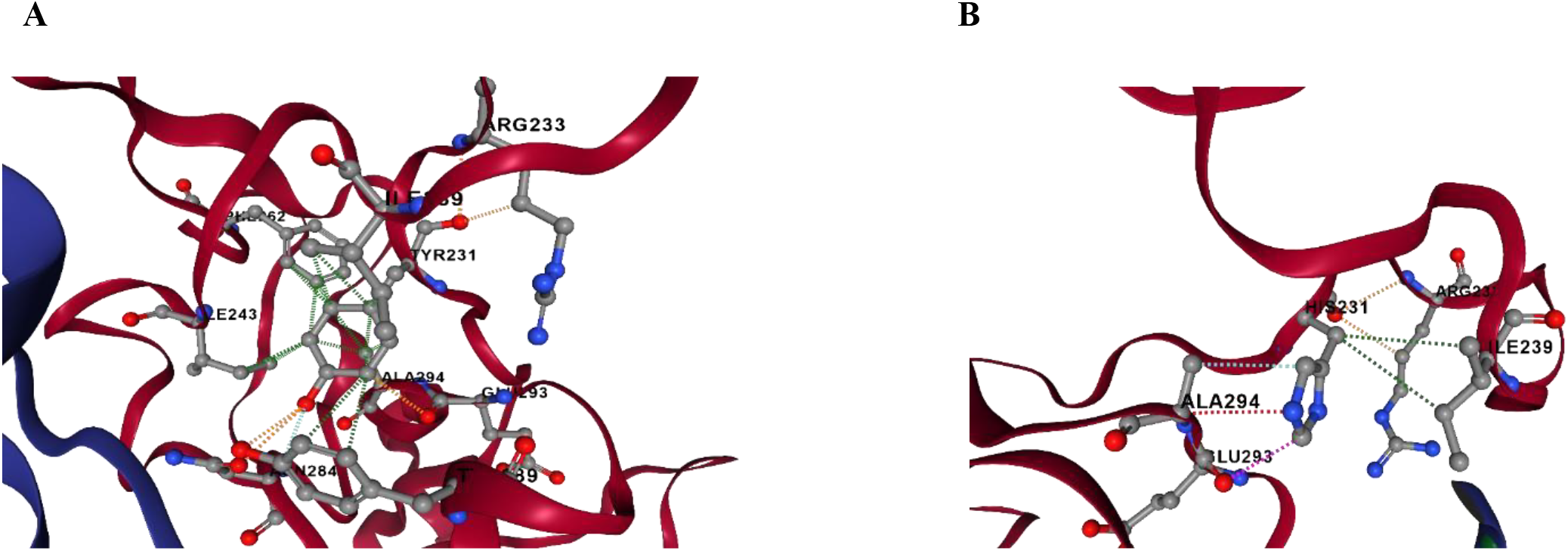
Structural comparison of the wild-type and p.Tyr231His ASPA protein predicted using DynaMut2. (A) Wild-type structure showing the native Tyr231 residue and its local interaction network. (B) Mutant structure showing the His231 substitution with reduced local residue interactions. The Y231H mutation was predicted to destabilize the protein, with a ΔΔG value of −2.04 kcal/mol.

To further evaluate the functional consequences of the ASPA p.Tyr231His variant, multiple in silico pathogenicity prediction tools were employed. Computational analyses using MutationTaster, SIFT, CADD, REVEL, and PolyPhen2 consistently predicted the variant to be deleterious or pathogenic, providing concordant evidence for its damaging functional effect (Table 1).

**Table 1.** In silico functional prediction of the ASPA variant using multiple computational tools.

| Tool | Score | Interpretation |
| --- | --- | --- |
| CADD | 25.3 | Predicted deleterious |
| REVEL | 0.913 | Likely pathogenic |
| PolyPhen-2 | 0.979 | Probably damaging |
| MutationTaster | - | Disease-causing |
| SIFT | 0.00 | Damaging |

The variant achieved a CADD score of 25.3 and a REVEL score of 0.913, both indicative of a high likelihood of pathogenicity. Similarly, PolyPhen-2 predicted the substitution to be probably damaging (score = 0.979), while SIFT classified the variant as damaging (score = 0.00). MutationTaster further identified the mutation as disease-causing. The concordance observed among multiple independent prediction algorithms provides strong computational evidence supporting the pathogenic nature of the ASPA p.Tyr231His variant and suggests that the substitution is likely to impair normal ASPA protein function.

### 3.2 Evolutionary conservation analysis of the ASPA p.Tyr231His variant

Multiple sequence alignment of ASPA orthologs from Homo sapiens, Mus musculus, Rattus norvegicus, Xenopus tropicalis, and Danio rerio using Clustal Omega demonstrated that Tyr231 is completely conserved across all analyzed vertebrate species (Fig. 3). The high degree of evolutionary conservation indicates that Tyr231 is under strong evolutionary constraint and is likely critical for maintaining the structural integrity and functional maintenance of ASPA. Consequently, substitution of this highly conserved tyrosine residue with histidine (p.Tyr231His) may alter the local protein environment and contribute to reduced structural stability, consistent with the DynaMut2-based stability prediction.

**Figure 3.**
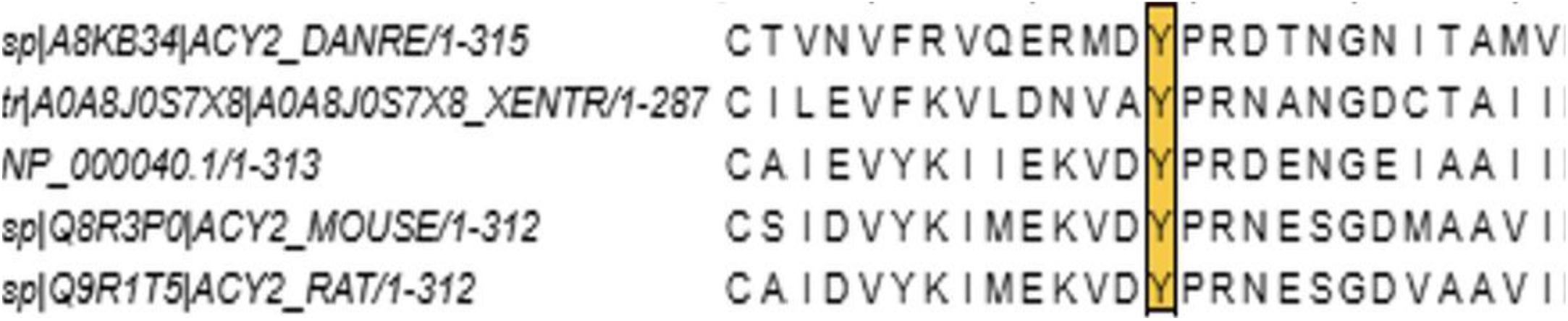
Multiple sequence alignment of ASPA orthologs from human, mouse, rat, Xenopus, and zebrafish. The Tyr231 residue (highlighted) is completely conserved across all analyzed vertebrate species, indicating strong evolutionary conservation and suggesting a potential structural and functional role of this residue.

### 3.3 Design and optimization of sgRNA for CRISPR-based base editing

Candidate sgRNAs targeting the ASPA c.691T>C (p.Tyr231His) variant were identified and comparatively evaluated based on editing precision, predicted bystander editing, PAM compatibility, GC content, MIT specificity score, CFD score, and structural suitability. Both candidate sgRNAs positioned the pathogenic cytosine within the optimal editing window of the cytosine base editor and were adjacent to canonical NGG PAM sequences, supporting compatibility with Cas9 targeting requirements.

Comparative analysis revealed that although both sgRNAs positioned the pathogenic cytosine within the editing window, sgRNA-1 was predicted to enable precise correction without introducing additional editable cytosines or predicted bystander alterations. In contrast, sgRNA-2 contained additional cytosines within the editing window, increasing the likelihood of unintended coding changes. Therefore, sgRNA-1 was selected for downstream analyses (Table 2).

**Table 2.**
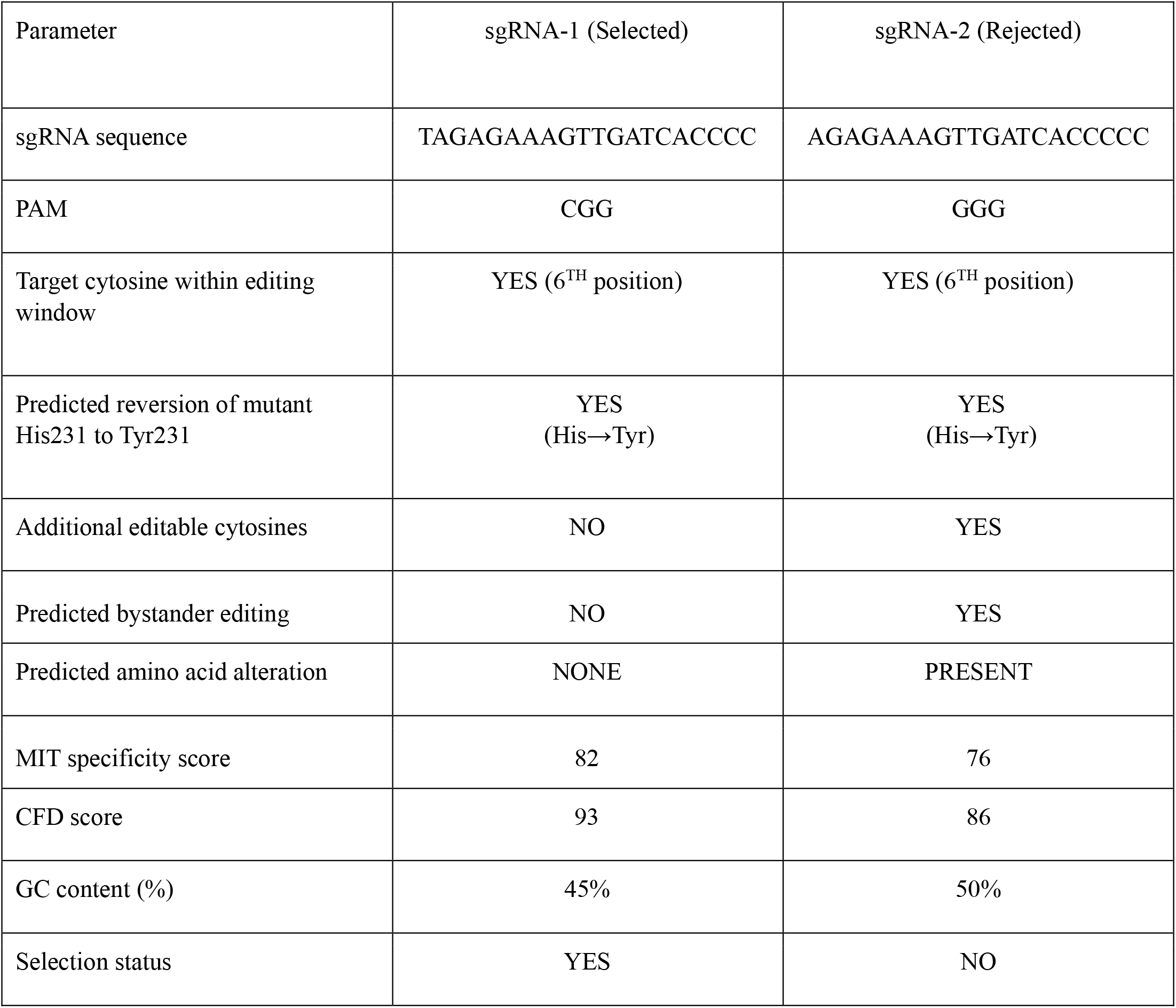
Comparative assessment of candidate sgRNAs for ASPA base editing.

Secondary structure prediction using RNAfold revealed that the selected sgRNA adopts a predominantly unstructured conformation, as indicated by the absence of predicted intramolecular base pairing within the minimum free energy (MFE) structure, as shown in Fig. 4. The RNAfold-predicted secondary structure consisted entirely of unpaired nucleotides, suggesting high structural accessibility and minimal thermodynamic constraints on sgRNA–target hybridization.

**Fig 4.**
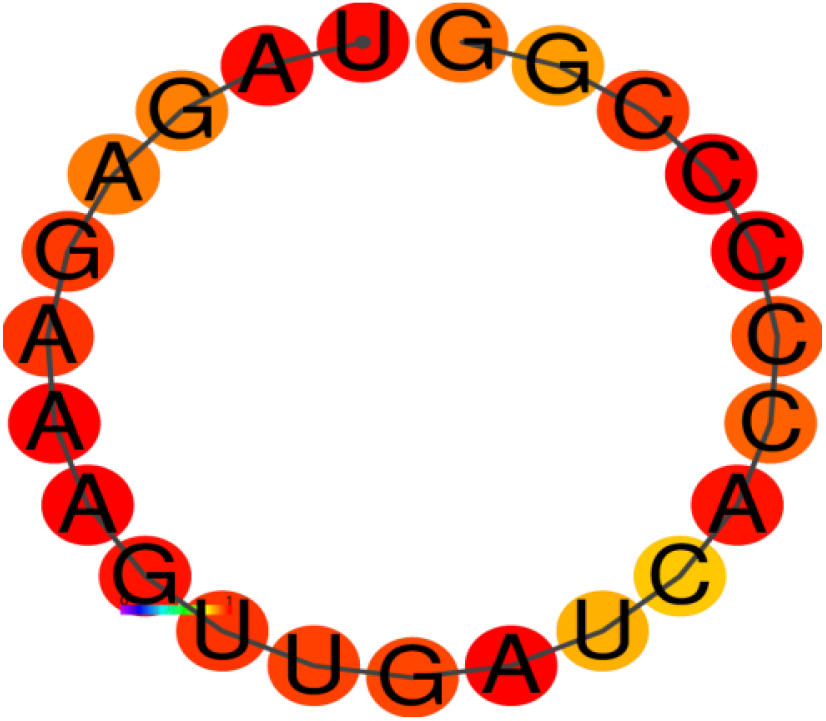
RNAfold-predicted secondary structure of the selected sgRNA. The guide RNA adopts a predominantly unstructured conformation with no predicted internal base pairing and a minimum free energy (MFE) of 0.00 kcal/mol, indicating favourable predicted accessibility for target recognition.

The predicted MFE was 0.00 kcal/mol, suggesting negligible thermodynamic constraints on guide folding. Similarly, the free energy of the thermodynamic ensemble was only −0.22 kcal/mol, indicating minimal secondary structure formation across the ensemble of possible conformations. The MFE structure accounted for 70.12% of the thermodynamic ensemble, indicating that a single dominant conformation is strongly favoured. Furthermore, the low ensemble diversity value of 1.13 indicates limited structural variability among predicted conformations. Collectively, these findings suggest that the sgRNA remains highly accessible for target recognition and Cas9 complex formation, supporting its potential suitability for CRISPR-mediated editing.

### 3.4 Genome-wide off-target profiling and specificity analysis of the designed sgRNA

To evaluate the specificity of the designed sgRNA, a comprehensive genome-wide off-target analysis was performed using CRISPRitz. The analysis identified a limited number of potential off-target sites across the human genome, indicating a favourable specificity profile. A total of 108 potential off-target sites were identified. No off-targets containing one or two mismatches were detected, whereas seven sites contained three mismatches and 101 sites contained four mismatches, as shown in Fig. 5. The absence of low-mismatch off-target candidates suggests strong sequence discrimination by the sgRNA and a reduced likelihood of unintended targeting events. A single perfect-match site corresponding to the intended ASPA target locus was identified, indicating the presence of a unique perfect-match site corresponding to the intended ASPA target locus within the analysed genome. Furthermore, bulge-aware off-target analysis was performed to identify potential DNA or RNA bulge-mediated recognition events.

**Fig 5.**
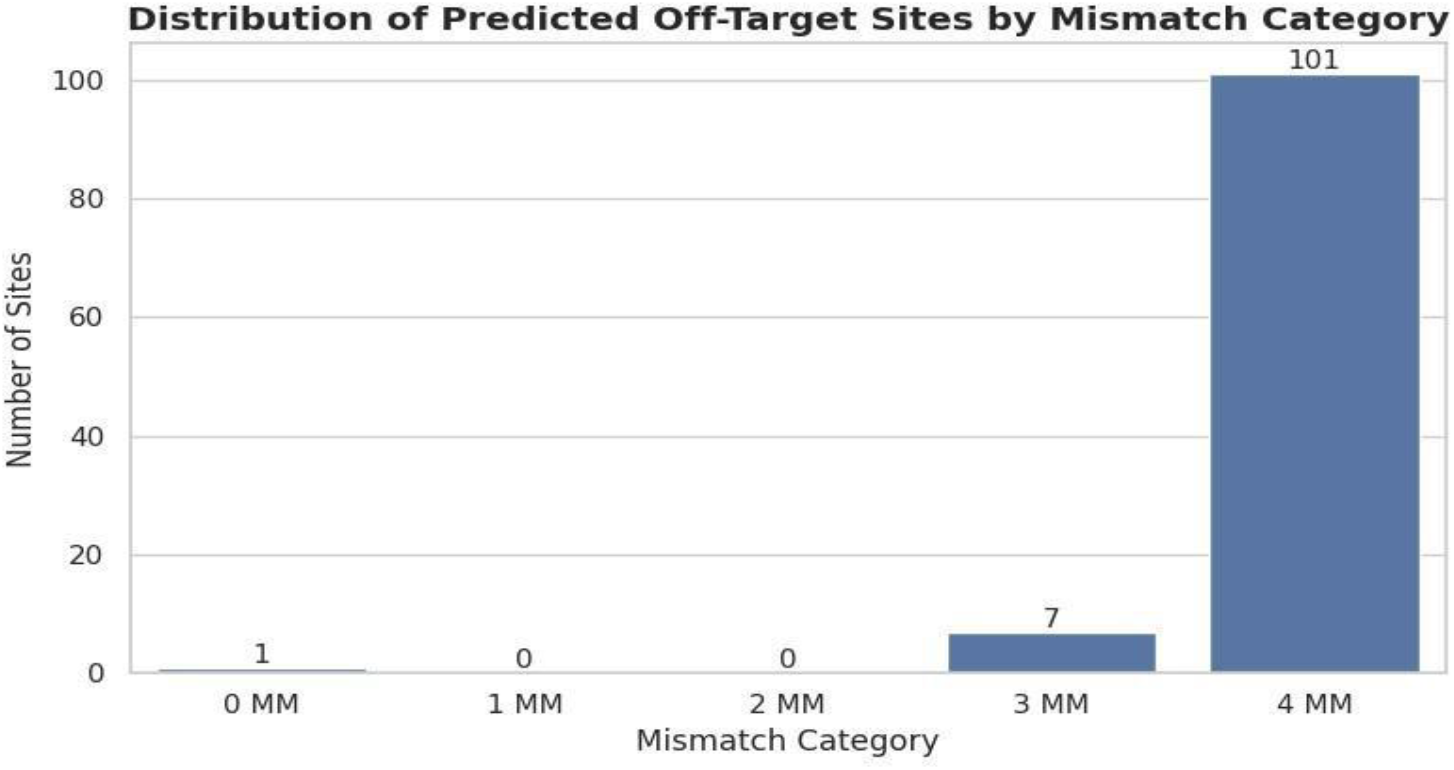
Mismatch distribution of genome-wide target sites identified by CRISPRitz. One perfect-match site (0 mismatches) corresponded to the intended on -target locus, whereas all predicted off -target sites contained >3 mismatches. Among the 108 off target sites identified, 7 exhibited three mismatches, and 101 exhibited four mismatches, indicating high predicted sgRNA specificity.

No off-target sites containing DNA or RNA bulges were detected, as all identified candidate sites exhibited a bulge size of zero. These findings further support the predicted high sequence specificity of the selected sgRNA and indicate limited tolerance for non-canonical guide–target interactions.

To investigate the positional distribution of mismatches across the guide sequence, a positional mismatch analysis was performed. As shown in Fig. 6, mismatches were not evenly distributed across the 20-nucleotide sgRNA. Certain positions, particularly BP12, BP13, BP15, and BP17, exhibited higher cumulative mismatch frequencies, whereas positions such as BP4 and BP6 showed relatively few mismatches, suggesting that these sites are more conserved during target recognition. Most mismatch events were associated with predicted off-targets containing four mismatches, while only a small number occurred among three-mismatch off-targets. This indicates that substantial sequence divergence is generally required before alternative genomic sites become potential off-targets, supporting the high specificity of the selected guide RNA. It is important to note that the heatmap represents the cumulative number of mismatches observed at each guide position across all predicted off-target alignments rather than the total number of off-target sites. Overall, the positional mismatch profile suggests that the selected sgRNA has strong predicted sequence specificity while exhibiting limited tolerance to sequence variation at key guide positions.

**Fig. 6.**
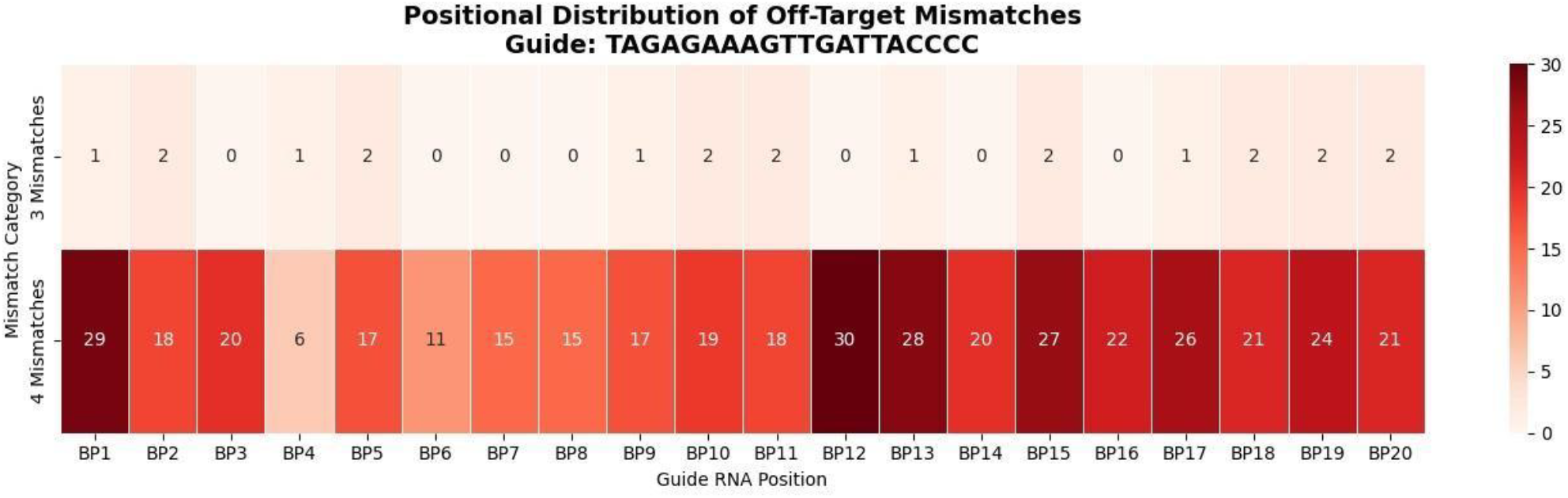
Position-specific distribution of mismatches across the selected sgRNA. The heatmap shows the cumulative frequency of mismatches observed at each nucleotide position (BP1–BP20) among predicted off-target sites identified by CRISPRitz. Darker colours indicate higher mismatch frequencies. The majority of mismatch events were associated with off-target sites containing four mismatches, whereas relatively few occurred among three-mismatch sites, indicating that substantial sequence divergence is generally required for off-target recognition.

Genomic annotation of the predicted off-target sites showed that most were located within intronic regions (n = 62), followed by DNase hypersensitive regions (n = 27). In contrast, only three off-target sites mapped to exonic sequences, suggesting a low probability of directly disrupting protein-coding genes. The remaining off-targets were distributed across regulatory genomic elements, including promoter-flanking regions, enhancers, transcription factor binding sites, and CTCF-binding sites, although each accounted for relatively few predicted sites. Overall, the off-target profile was predominantly associated with non-coding genomic regions, suggesting a reduced likelihood of directly affecting protein-coding regions. The distribution of predicted off-target sites across different genomic features is presented in Figure 7.

**Fig 7.**
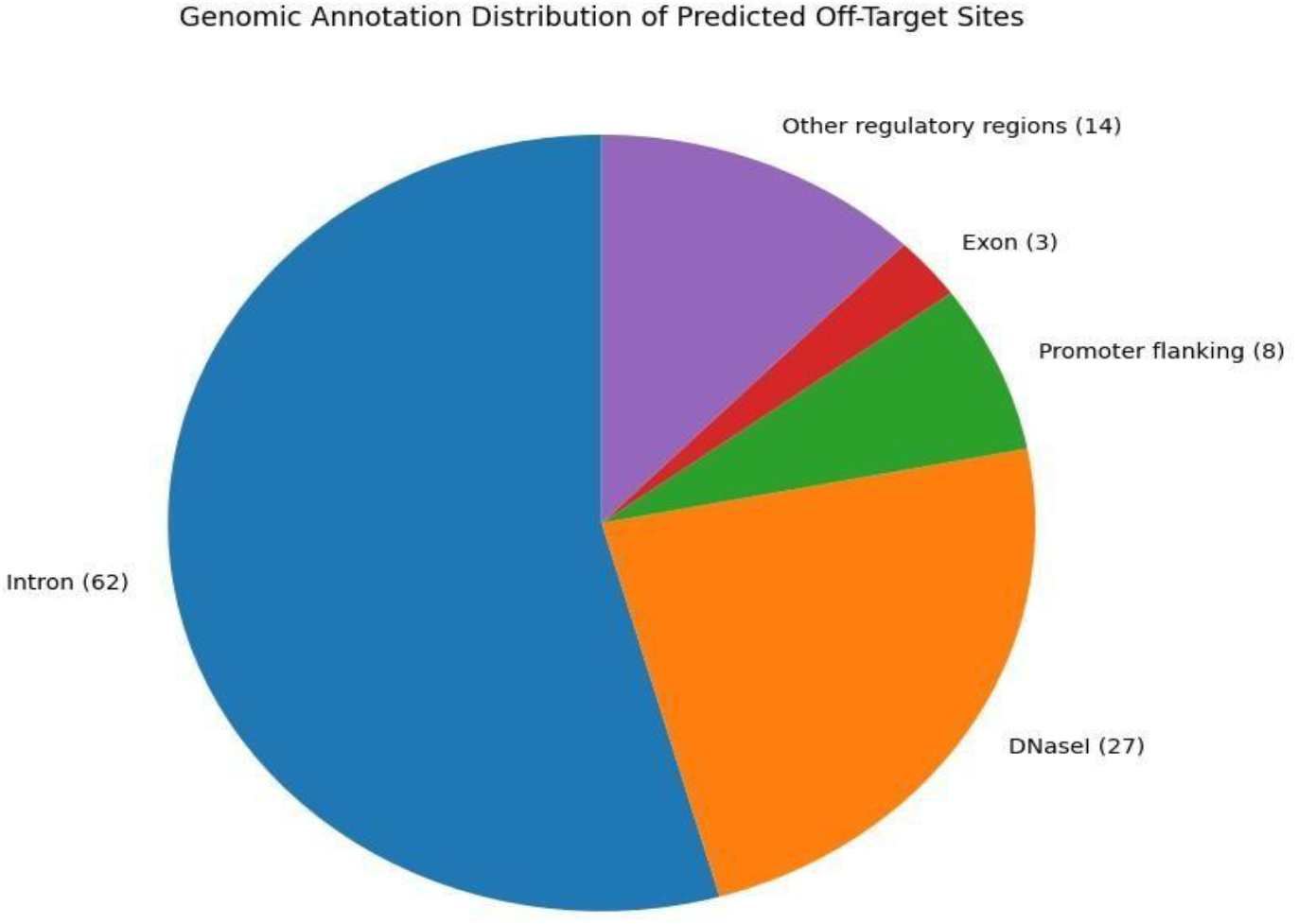
Genomic annotation profile of predicted off-target sites. Distribution of predicted off-target loci among genomic features, including introns, exons, promoters, enhancers, DNase I hypersensitive regions, transcription factor binding sites, and other regulatory elements. Intronic regions constituted the largest proportion of predicted off-targets, whereas exonic sites were comparatively rare.

## 4. Discussion

The rapid development of CRISPR–Cas genome editing technologies, together with advances in high-throughput sequencing, has greatly improved the identification and correction of disease-associated genetic variants (Kim et al.

2024). In the present study, an integrated in-silico workflow was used to design and evaluate a cytosine base editing (CBE) strategy targeting a pathogenic mutation in the *ASPA* gene. Unlike conventional CRISPR–Cas9 systems that rely on double-strand DNA breaks, cytosine base editors enable precise C-to-T nucleotide conversion without generating double-strand breaks, thereby reducing the likelihood of indel formation associated with DNA cleavagebased editing approaches.

The designed sgRNA positioned the target cytosine within the canonical editing window (positions 4–8 relative to the PAM), which has been reported to favour efficient cytosine base editing in previous studies. The sgRNA also demonstrated favourable design characteristics, including balanced GC content and high specificity scores, both of which are recognised contributors to efficient target recognition and reduced off-target activity. Furthermore, RNAfold analysis predicted a relatively unconstrained secondary structure with limited intramolecular base pairing. Previous studies have shown that excessive secondary structure can hinder guide accessibility and reduce Cas9 loading efficiency; therefore, the predicted folding pattern may be favourable for guide accessibility and target recognition.

Genome-wide off-target analysis further demonstrated a high level of predicted specificity, with no potential off-target sites identified below three mismatches. Most predicted off-targets contained four mismatches and were predominantly located within intronic or other non-coding genomic regions. This finding is encouraging because off-target events occurring within coding exons may have a greater potential to affect gene function compared with those located in non-coding regions. However, regulatory elements can also influence gene expression. Consequently, the observed genomic distribution suggests a lower likelihood of predicted off-target effects within protein-coding regions; however, experimental validation remains necessary to determine the biological significance of these sites. These findings also highlight the value of integrating complementary computational tools, including CRISPOR, CRISPRitz and RNAfold, to improve confidence in sgRNA selection before experimental validation.

Despite these promising findings, the present study is limited to computational predictions and therefore requires experimental validation. The editing efficiency, editing purity, and potential bystander editing predicted in silico may differ under biological conditions. In addition, chromatin accessibility, cellular DNA repair mechanisms, base editor expression levels, editor-dependent activity, and delivery strategies may influence editing outcomes in vivo and are not captured by the computational analyses performed in this study. Future investigations should validate the proposed sgRNA in relevant cell-based and animal models to assess editing efficiency, specificity, long-term genomic safety, and therapeutic feasibility.

Overall, this study provides a comprehensive computational framework for the rational design and evaluation of a cytosine base editing strategy targeting the ASPA c.691T>C (p.Tyr231His) variant associated with Canavan disease. By integrating variant characterization, structural assessment, sgRNA optimization, secondary structure analysis, and genome-wide off-target profiling, the proposed workflow identifies a highly specific candidate guide with favourable predicted editing characteristics. Although experimental validation remains essential to confirm editing efficiency, specificity, and functional restoration of ASPA activity, these findings demonstrate the potential of in silico approaches to accelerate the development of precision genome-editing strategies for rare monogenic disorders. This study establishes a foundation for future experimental investigations and highlights the growing role of computational pipelines in guiding safer and more targeted therapeutic genome engineering approaches.

## CRediT authorship contribution statement

Aditi Verma: Conceptualization, Methodology, Formal analysis, Visualization, Writing – original draft. Elamathi Natarajan: Supervision, Methodology, Review & Editing.

## Data availability

The datasets analyzed during the current study are publicly available from NCBI RefSeq, ClinVar, gnomAD, UniProt, Protein Data Bank (PDB), and the web servers cited in the manuscript. No new datasets were generated during this study.

## Ethics statement

This study was conducted entirely using publicly available genomic and protein databases and computational bioinformatics tools. No human participants, animals, or identifiable patient data were involved. Therefore, ethical approval and informed consent were not required.

## Declaration of Generative AI and AI-assisted technologies in the writing process

During the preparation of this manuscript, the authors used generative artificial intelligence (AI)-based tools to assist with language refinement and improvement of manuscript clarity. The authors have reviewed and edited all AI-assisted content and take full responsibility for the accuracy, integrity, and originality of the published work.

